# Origin of Neural Frequency Responses: Intrinsic Sensory Coding vs. Structural Influences

**DOI:** 10.1101/2023.10.17.562345

**Authors:** Ilker Duymaz, Naoki Kogo, Nihan Alp

**Author notes:** Correspondence concerning this article should be addressed to Ilker Duymaz, Mathematisches Institut, Justus-Liebig-Universität Gießen, Arndtstraße 2, 35392 Gießen.

## Abstract

Periodic changes in visual input can produce rhythmic patterns in EEG signals, which appear as narrowband frequency components. These components are commonly interpreted as reflecting the activity of neurons sensitive to the modulated stimulus features. Here, we present a scenario in which frequency components arise solely from retinotopic variations in signal strength, without reflecting any specific neural mechanism sensitive to the modulated feature. Using simulated and empirical data, we show that signal fluctuations based purely on retinotopic stimulus position can produce identifiable frequency components in response to position-modulated stimuli. These components likely reflect structural rather than functional cortical factors influencing signal strength across different retinotopic areas. Our results challenge the conventional assumption that frequency components necessarily indicate intrinsic neural signal processing, instead highlighting how interactions between stimuli and cortical architecture can give rise to such components.

## Introduction

Rhythmic changes in visual stimuli can induce corresponding rhythmic patterns in brain activity, which appear in EEG recordings as narrowband frequency components. When a visual or semantic stimulus property (e.g., contrast, object identity) is modulated at a specific frequency, neurons sensitive to that property tend to oscillate at the same or related frequencies, resulting in periodic fluctuations in EEG amplitude. By presenting the stimulus for a sufficient duration (e.g., longer than one second; Norcia et al. (2015)) and applying frequency-domain analyses, these periodic fluctuations can be identified as precise, narrowband frequency components. This phenomenon, known as the steady-state visually evoked potential (SSVEP; Regan (1977)), allows researchers to extract stimulus-related brain activity with a high signal-to-noise ratio. As a result, SSVEPs have become a powerful tool for probing visual processing at multiple levels (Allen et al., 1996; Alp & Ozkan, 2022; Alp et al., 2016,2018; Gundlach & Müller, 2013; Rossion et al., 2015), characterizing visual and neurological impairments (Sartucci et al., 2010; Silberstein et al., 2000), and developing brain-computer interface (BCI) technologies (see Liu et al. (2022) for a review).

The distinctive inferential power of SSVEPs is rooted in the presumption that the precise frequency components evident in the data can be attributed to modulations in neuronal activity, particularly of cells sensitive to the modulated stimulus property. In theory, any other factor capable of inducing fluctuations in EEG amplitude could also give rise to such frequency components. However, for these fluctuations to be practically relevant, they must withstand the standard practices in experimental design and data processing that are typically used to filter out extraneous signals.

Of particular significance of these practices are extended trial durations and the averaging of data across multiple trials, which serve to accentuate visually evoked potentials by removing transient fluctuations that are not time-locked across trials (Trimble, 1968). Consequently, induced fluctuations must exhibit consistency within trials over time and synchronized phase across multiple trials to produce identifiable frequency components. Furthermore, given that SSVEP studies often focus exclusively on frequency components related to the modulation frequency of a periodically modulated stimulus property (Norcia et al., 2015), these fluctuations must also display periodic components that correspond to the stimulus modulation frequency and its harmonics.

Admittedly, it is challenging to envision any other factors capable of generating highly consistent signal fluctuations that are also sensitive to modulations in visual stimuli. Particularly, the latter condition suggests a level of computational capability to process the modulated aspects of stimuli. Given the extensive array of low (e.g., contrast, motion direction) and high-level (e.g., face identity, object category) stimulus properties that can produce frequency components when modulated, it is clear that neuronal responses tuned to specific stimulus properties are the primary driver of these signal fluctuations. Consequently, alternative sources for these fluctuations have traditionally been overlooked, as no compelling scenario has emerged in which any other factor could consistently introduce signal fluctuations that align with the modulations in visual stimuli.

Nonetheless, the magnitude of neuronal activity is not the only factor that determines the strength of measured EEG signals. Various other factors, such as the cortical volume (Schaul, 1998) and depth of the activated regions (Butler et al., 2019), the signal cancellation between simultaneously active brain areas (Ahlfors et al., 2010), and the thickness of the cerebrospinal fluid layer covering regions of interest (Rice et al., 2013), can impact the translation of cell activity into EEG signals. Under certain circumstances, such structural factors, as opposed to functional, can potentially interact with stimuli to introduce superficial influences into the data by modulating EEG amplitudes across conditions and/or time.

Particularly, position-dependent factors that introduce variances in the properties of different retinotopic populations can have a potentially large impact on recorded EEG signals by interacting with stimulus position. The visual field is not uniformly mapped onto the visual cortex (Allman & Kaas, 1971; Daniel & Whitteridge, 1961; Talbot & Marshall, 1941; Wandell et al., 2007). Significant differences exist in cortical volume and depth across retinotopic areas, depending on eccentricity and polar angle (Daniel & Whitteridge, 1961; Horton & Hoyt, 1991; Schwartz, 1980). When a stimulus is located in the periphery, a given portion of the visual field engages smaller and deeper neural populations (Daniel & Whitteridge, 1961; Horton & Hoyt, 1991; Schwartz, 1980).

Similarly, population receptive field sizes are much smaller in foveal areas compared to peripheral ones, meaning that a central stimulus would engage a greater volume of neural tissue (Dumoulin & Wandell, 2008). As a result, the same stimulus can evoke larger or smaller EEG responses depending on the size and depth of the activated brain areas based on its position in the visual field (Slotnick et al., 2001).

For SSVEP studies in general, the modulation of EEG signals by such position-dependent factors does not directly warrant identifiable frequency components in the signal. In order to yield frequency components, these position-dependent factors would somehow have to induce consistent and phase-locked signal fluctuations across many repetitions of a condition. When stimuli are static, or when they do not follow a consistent pattern of motion across trials, it is highly unlikely for position-dependent factors to induce signal fluctuations that survive trial-averaging. However, if the position of a stimulus is periodically modulated in a consistent manner across trials, as is often the case in SSVEP studies involving moving stimuli (e.g., Aissani et al., 2011; Nozaradan et al., 2012; Pitchaimuthu et al., 2021; Varlet et al., 2023), these position-dependent imbalances may introduce consistent time-locked fluctuations in the signal, potentially giving rise to narrowband frequency components even after averaging signals across trials (Figure 1).

**Figure 1.**
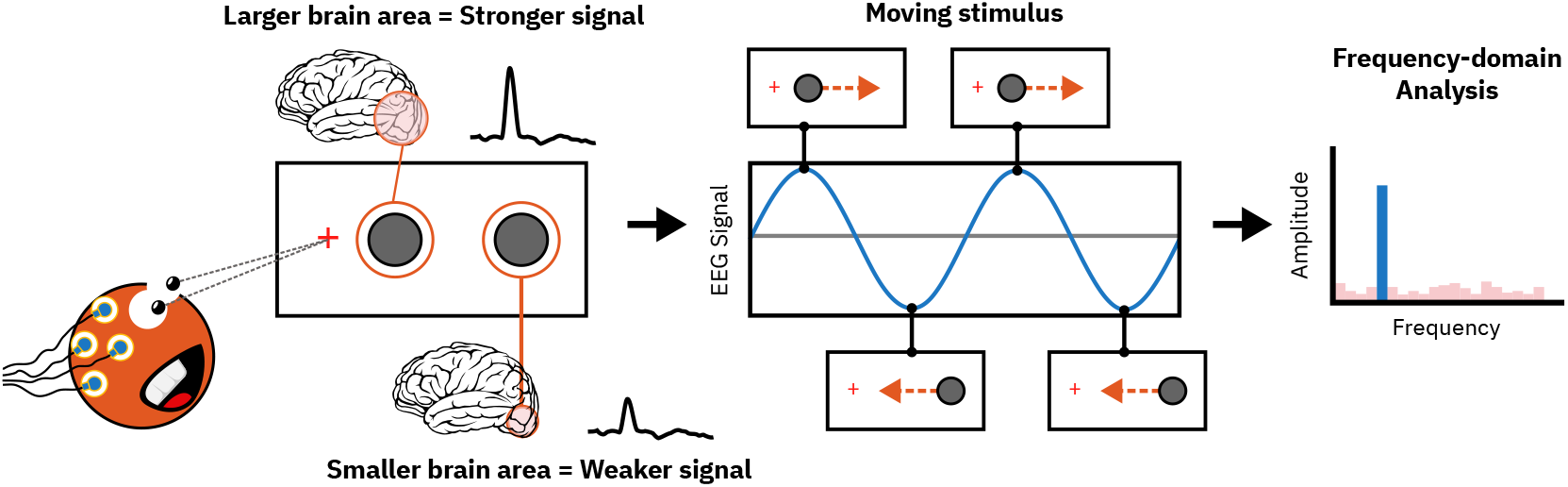
Frequency components arising from position-dependent periodic fluctuations. This figure depicts a hypothetical situation based on the known eccentricity-related variations in cortical magnification (Daniel & Whitteridge, 1961; Horton & Hoyt, 1991; Schwartz, 1980). If the brain allocated equal-sized and equally-dense cortical regions to different parts of the visual field, a singleton moving stimulus would generate a uniform, consistent neural response across all positions, resulting in a stable, unvarying signal (gray line). On the other hand, if different areas in the visual field are represented disproportionately in the visual cortex, the same stimulus can lead to stronger or weaker signals depending on its position and the size of the corresponding brain regions (blue line). When the stimulus position is periodically modulated, this could lead to the emergence of periodic fluctuations in the EEG signal, which could yield identifiable frequency components.

In this context, we present a scenario in which stimulus-related frequency components emerge exclusively due to retinotopic variations in signal strength, rather than as a result of neural processing by specific feature-tuned neurons. First, we use simulations to demonstrate that any sort of position-dependent variance in measured EEG responses can potentially introduce periodic signal fluctuations by interacting with position-modulated stimuli even when the responses of local receptive units are driven equally. If these fluctuations are discernible in actual EEG signals, they should manifest even in the absence of modulations in other stimulus properties, and must be strictly phase-coupled to modulations in stimulus position. In two subsequent EEG experiments, we demonstrate that periodically modulating the polar angle and eccentricity of a singleton stimulus induces frequency components when all other stimulus properties remain constant over time. Critically, these frequency components are eliminated through phase-cancelation when the phases of the position modulations are varied across trials.

If these frequency components were generated solely by the intrinsic neural processing of the stimulus, they would have persisted even in the phase-varied conditions. Phase-cancelation in the phase-varied conditions suggests that these frequency components are phase-coupled to the position modulation of the stimulus, and therefore must strictly be related to fluctuations produced by position-dependent factors. Taken together, these results indicate that inherent retinotopic differences in measured EEG amplitude can autonomously introduce consistent, time-locked signal fluctuations in response to position-modulated stimuli, offering a counterexample to the conventional approach of associating frequency components with intrinsic neural signal processing.

### Simulation

It is conceivable to view the EEG signal as a comprehensive non-linear transformation on visual input. This transformation is the culmination of numerous layers of diverse non-linear processes introduced by the intricate machinery of neural processing, in addition to the physical factors that govern the translation of neuronal activity into EEG signals. For our purposes, we draw a clear distinction between two pivotal concepts: unit response and population-level response. Following this distinction, we will argue that while traditional explanations attribute stimulus-related frequency components to the responses of units tuned to the modulated stimulus properties, these components can also emerge at the level of population response. This emergence hinges on variances in the population response contingent upon the retinotopic stimulus position and the presence of a position-modulated stimulus.

We start by conceptualizing the unit response as an abstract representation of neuronal responses triggered by visual stimuli. In this context, the unit response characterizes a transformation of visual input, which we define as an arbitrary nonlinear function, denoted as *y* = *f* (*xs*(*t*), *a*(*t*)). Here, *y* signifies the response amplitude of a unit responding to a stimulus whose position is *xs* and intensity *a* at time *t*. Our focus is not on modeling specific types of neurons, but rather on defining units that adjust their responses based on varying intensities of the stimulus properties to which they are tuned. Therefore, *a* (i.e., stimulus intensity) can be associated with any stimulus property to which these hypothetical units are sensitive, and the function *f* can encompass various forms of neural computations.

The conventional interpretation of the frequency components observed in EEG signals in response to a periodically modulated stimulus assumes that these frequency components arise from the nonlinear relationship between *a* and *y*. As long as *f* includes a non-zero coefficient for *a*, indicating sensitivity to changes in stimulus intensity, periodic modulation of *a* will invariably result in a periodic modulation of *y*. Consequently, if the stimulus intensity undergoes sinusoidal modulation, for example, it will induce a non-sinusoidal modulation of the unit response that repeats at the same time intervals as the stimulus modulation. As a result, the EEG signal will contain frequency components that are harmonics (integer multiples) of the sinusoidal stimulus modulation frequency, provided that an adequate number of units synchronously respond to the stimulus.

Here, our aim is to investigate whether identifiable frequency components can be generated independently of the unit response, specifically due to position-dependent variances in the responses of different retinotopic populations. Therefore, we focus on the scenario in which the stimulus intensity α remains constant across time and the unit responses do not vary as a function of modulations in stimulus intensity. If a position-modulated stimulus, across its different positions, was processed by a perfectly homogenous ensemble of identical units without any position-dependent variability in the measured EEG strength, the resulting signal would not exhibit fluctuations and remain constant across time. This is because the stimulus would drive uniform responses from the same number of units with identical properties at each time point (Figure 1, gray line). Conversely, if there are position-dependent factors that influence how unit responses translate into measured EEG strength, such as the density of units or their distance from the electrodes, the signal would exhibit fluctuations across different positions of the stimulus, even though individual local units receive the same information (i.e., stimulus intensity) and produce the same response. Moreover, if there are consistent variations in the receptive field properties of different retinotopic populations, these could also lead to position-dependent variability in EEG signals by modulating unit responses based on the pattern of overlap between the receptive fields and stimulus (Figure 1, blue line).

To capture the effect of such position-dependent factors, we conceptualize population response as a representation of collective activity from many individual units responding to the stimulus as measured by the EEG. At each position of the stimulus, a number of units whose receptive fields overlap with the stimulus produce a response. The population response is a sum of responses from each of these units with weights determined as a function of the location of the unit’s receptive field within the visual field:

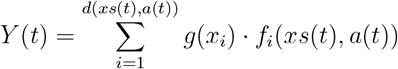

where *d* is a unit-density function that determines the number of units responding to a stimulus presented at retinotopic position *xs* with intensity *a* at time *t*; *f* is the magnitude of response for each unit responding to that stimulus; and *g* determines the extent to which the response of each unit contributes to the measured population response based on the location of its receptive field, *x*_*i*_. Therefore, *Y* is the magnitude of the measured population response evoked by a stimulus whose position is *xs* and intensity *a*.

As our focus is solely on signal variations arising from position-specific factors, we assume that individual units produce identical responses when a stimulus with the same intensity *a* falls within their receptive fields. Therefore, we factor out the variance attributable to the correlation between individual unit responses and stimulus intensity under the condition that the intensity *a* is constant. However, unit responses would still vary depending on the extent of overlap between each unit’s receptive field and the stimulus. Consequently, unit responses would exhibit variability contingent upon the positioning and dimensions of both their receptive fields and the stimulus. This inter-unit variance alone would not lead to differences in population response if the spatial distribution of units’ receptive field sizes were identical for each population. Conversely, if the receptive field sizes of individual units, or their spatial distribution within the population receptive field, vary across populations, this could lead to differences in population responses along with other position-dependent factors. We account for this by keeping the *f*_*i*_(*xs*(*t*)) term while excluding the variance attributable to *a*. Thus, we obtain:

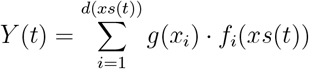

which denotes a stimulus-position-dependent coefficient for the magnitude of EEG signals recorded from a population of stimulated units, whose responses do not vary as a function of stimulus intensity but only as a function of the position of their receptive fields. It is then possible to calculate these position-dependent coefficients for stimuli encompassing arbitrary portions of the visual space (such as pixels on an image) at discrete time points and map them across the visual field. This produces a two-dimensional heatmap representing the variance in EEG amplitude caused by position-dependent factors given otherwise uniform unit responses; which we refer to as Retinotopic Weight Function (RWF):

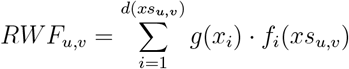

The actual nature of the Retinotopic Weight Function is not of critical importance for our demonstration, and it can be substituted by any two-dimensional function that may or may not approximate the influence of biologically-plausible position-dependent factors on the EEG strength. The critical requirement for the Retinotopic Weight Function in this context is that it consistently varies across the visual field and scales the contribution of each unit (to the signal) based on the location of their receptive fields. We will demonstrate that as long as this condition is fulfilled, position-dependent factors can interact with position-modulated stimuli to generate periodic signal fluctuations and therefore yield identifiable frequency components.

To this end, we simulate a hypothetical signal that contains only the fluctuations caused by position-dependent variances in population responses and not by the oscillations in unit responses induced by modulations in stimulus intensity. Therefore, we generate signals by multiplying a constant average unit response *C* with the respective position-dependent weight corresponding to the population response strength at each position of a moving stimulus.

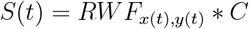

where *x* and *y* are the cartesian coordinates of the stimulus position.

Given this framework, we explore three different cases for the Retinotopic Weight Function, as depicted in Figure 2a-c. Although these Retinotopic Weight Functions were defined arbitrarily, some of them can be loosely associated with certain retinotopic properties of the visual cortex. For instance, in Figure 2a, the Retinotopic Weight Function is depicted as a function whose output diminishes linearly with distance from the center. It could be argued that eccentricity-dependent differences in cortical magnification could give rise to analogous variations in population response. In Figure 2b, the Retinotopic Weight Function exhibits a linear decrease from left visual field to right visual field and consequently introduces variations across polar angles, which could provide insights about the effect of retinotopic anisotropies. Finally, in Figure 2c, the outputs of the Retinotopic Weight Function are randomly determined from a uniform distribution in a fine grid to represent a complex spatially-varying function without assuming any specific computation. Although arbitrary, these Retinotopic Weight Functions all fulfill the essential condition of exhibiting consistent variations in measured population responses across different stimulus positions.

**Figure 2.**
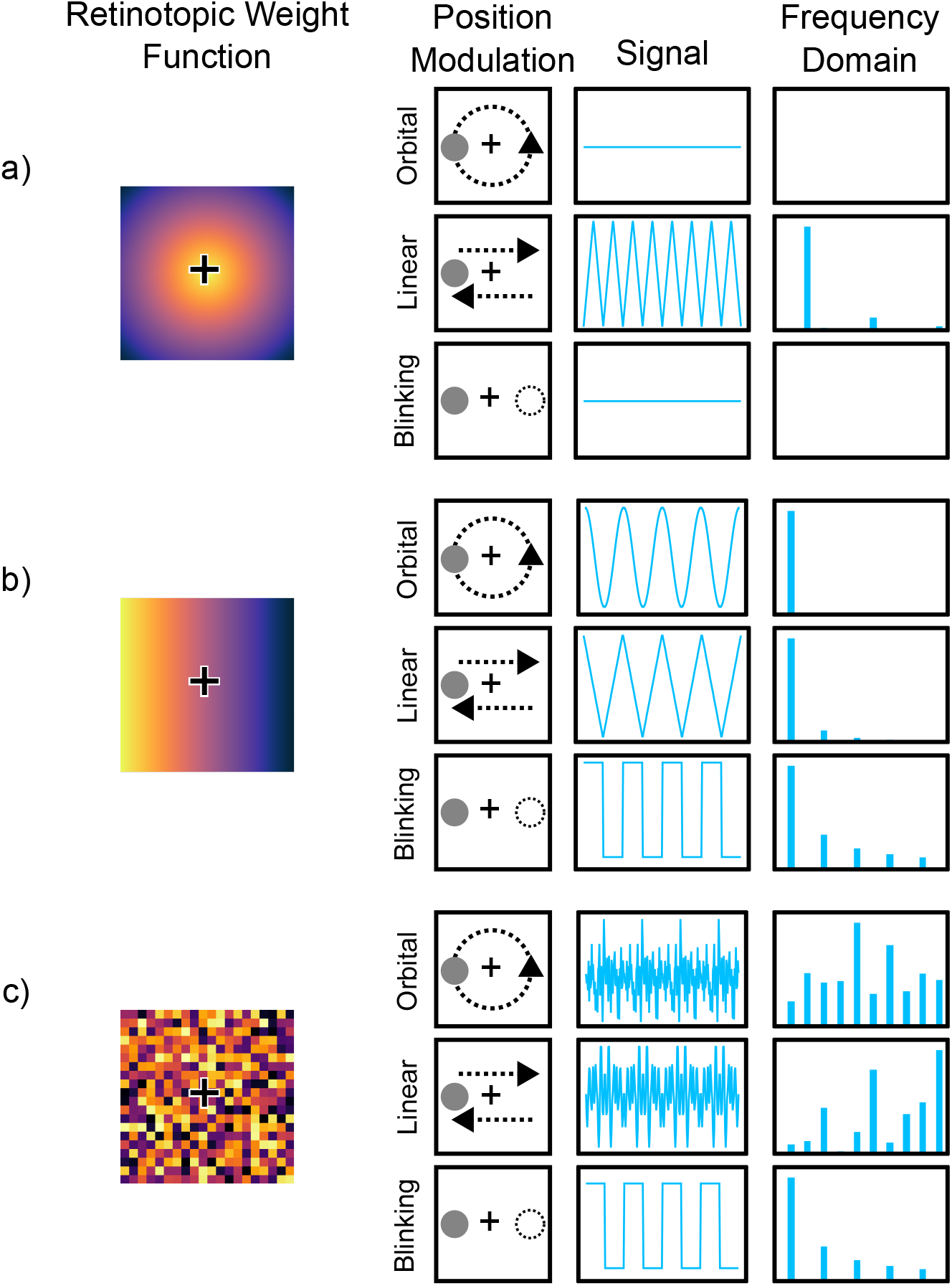
Simulated signals and their frequency domain representations. **Description for Figure 2:** The Retinotopic Weight Function (first column) interacts with the type of position modulation (second column: Orbital, Linear, Blinking) to shape the time domain response (third column) and resulting frequency content (last column). Frequency domain responses (last column) show harmonics of the position modulation frequency. **a)** Retinotopic Weight Function decreases linearly with eccentricity. **Orbital position modulation:** The singleton stimulus traverses along a circular path with constant eccentricity, which does not introduce fluctuations in the signal and does not lead to any frequency components. **Linear:** Stimulus eccentricity is linearly modulated along a horizontal line. The course of the eccentricity modulation is reversed sharply at the center and during motion-direction reversals (left-to-right, right-to-left). This is reflected in the signal as a triangular wave, which contains harmonics of double the position modulation frequency. **Blinking:** The singleton stimulus alternates between two locations that have identical weights in the Retinotopic Weight Function, therefore it also does not produce fluctuations and frequency components. **b)** Retinotopic Weight Function decreases linearly from left to right visual field. **Orbital:** The signal is the projection of a circular motion along a linear gradient, which by definition is sinusoidal. Therefore, only the fundamental position modulation frequency is represented in the frequency domain. **Linear:** Stimulus seesaws along the axis of linear Retinotopic Weight Function gradient. The signal exhibits sharp reversals when the stimulus changes direction. This triangular signal contains the odd harmonics of the position modulation frequency. **Blinking:** Stimulus alternates between two locations with different weights, leading to a square wave signal and harmonics of the position modulation frequency. **c)** Retinotopic Weight Function is randomly determined. **Orbital and linear position modulation:** The stimulus traverses through areas with randomly determined weights for the Retinotopic Weight Function. The consequent signals are highly complex, but since they have the same period as the position modulation cycle, they contain harmonics of the position modulation frequency in their frequency domain representations. **Blinking:** Stimulus alternates between two locations with different weights similarly to b). The square wave signal contains the harmonics of the position modulation frequency.

Finally, we examine various forms of position modulation applied to a singleton stimulus (Figure 2, Position Modulation column). First, we consider the rotational motion of a singleton object orbiting a central point (orbital). Second, we consider the repetitive translational motion of an object with a constant speed, which moves back and forth along a horizontal axis (linear). Third, we consider the position modulation of an object that blinks between two positions, akin to traditional long-range motion paradigms (blinking).

Figure 2 (Signal column) illustrates the simulated signals generated for all combinations of three distinct Retinotopic Weight Functions and three types of position modulation (i.e., orbital, linear, and blinking). Signals were generated for 4-second time windows with all three types of position modulation being repeated four times with a cycling frequency of 1 Hz. The simulated signals were then Fourier transformed to the frequency domain with a 1/4 = 0.25 Hz frequency resolution. All visible frequency components in the Frequency Domain column in Figure 2 correspond to the harmonics of the position modulation frequency (i.e., 1 Hz).

## Experiment 1: Polar angle modulation

### Materials and Methods

In Experiment 1, we investigated whether the modulated polar angle of a singleton stimulus could interact with position-dependent factors to produce distinct frequency components when other stimulus properties are kept constant.

#### Participants

14 Sabanci University students (8 female, 6 male; mean age = 21.1 years, SD = 2.02) with normal or corrected-to-normal vision participated in Experiment 1 for course credit. One participant’s data was excluded from all analyses due to software errors during the recording session, reducing our sample size to 13. The sample size was based on previous studies with similar methods (e.g., Nozaradan et al., 2012; Pitchaimuthu et al., 2021; Varlet et al., 2023). A sensitivity analysis conducted using GPower software (Faul et al., 2007) revealed that this sample size was sufficiently powered (power = .80) to detect large effect sizes (Cohen’s f > .42) for our within-subjects factors (see, Frequency domain analysis). All procedures in this study were reviewed and approved by the Sabancı University Research Ethics Council. Participants gave informed consent prior to the experiment session.

#### Apparatus and Stimuli

Stimuli were generated using Psychopy (Peirce et al., 2019) and presented on an ASUS XG248Q monitor (23.8”, resolution=1920×1080) with a refresh rate of 60 Hz. Participants viewed the stimuli from a 76.5 cm distance in a dark and quiet room. The stimuli consisted of a single light gray (contrast = 75%) dot rotating around a fixation cross on a dark gray (contrast = 25%) background. The dot had a radius of 1 dva and an eccentricity of 5 dva from the fixation cross. The dot always completed one full rotation in exactly 1 second and had a constant movement speed with a 6° polar angle shift per motion frame. Trials lasted 11 seconds, during which the dot completed 11 full rotations.

The rotational motion of the dot meant that its polar angle was modulated as a function of time. The phase of this polar angle modulation depended on the position from which the dot started its rotation. In one condition, the dot’s rotation always started from the same predetermined polar angle chosen randomly for each participant from a range of 0° to 350° in steps of 10° (e.g., 50° for one participant, 130° for another). The constant starting polar angle led to a phase-locked manipulation of the dot’s polar angle across trials (Figure 3a). We will hereafter refer to this condition as the phase-locked (PL) condition. In the second condition, the starting polar angle was randomly selected for each trial from the same range (0° to 350° in steps of 10°) without replacement (i.e., each step was used only once). Since the dot started its rotation from a different polar angle on each trial, the polar angle modulation was not phase-locked in this condition (Figure 3b). Therefore, we will refer to this condition as the phase-varied (PV) condition. There were 72 trials for each phase condition (PL and PV).

**Figure 3.**
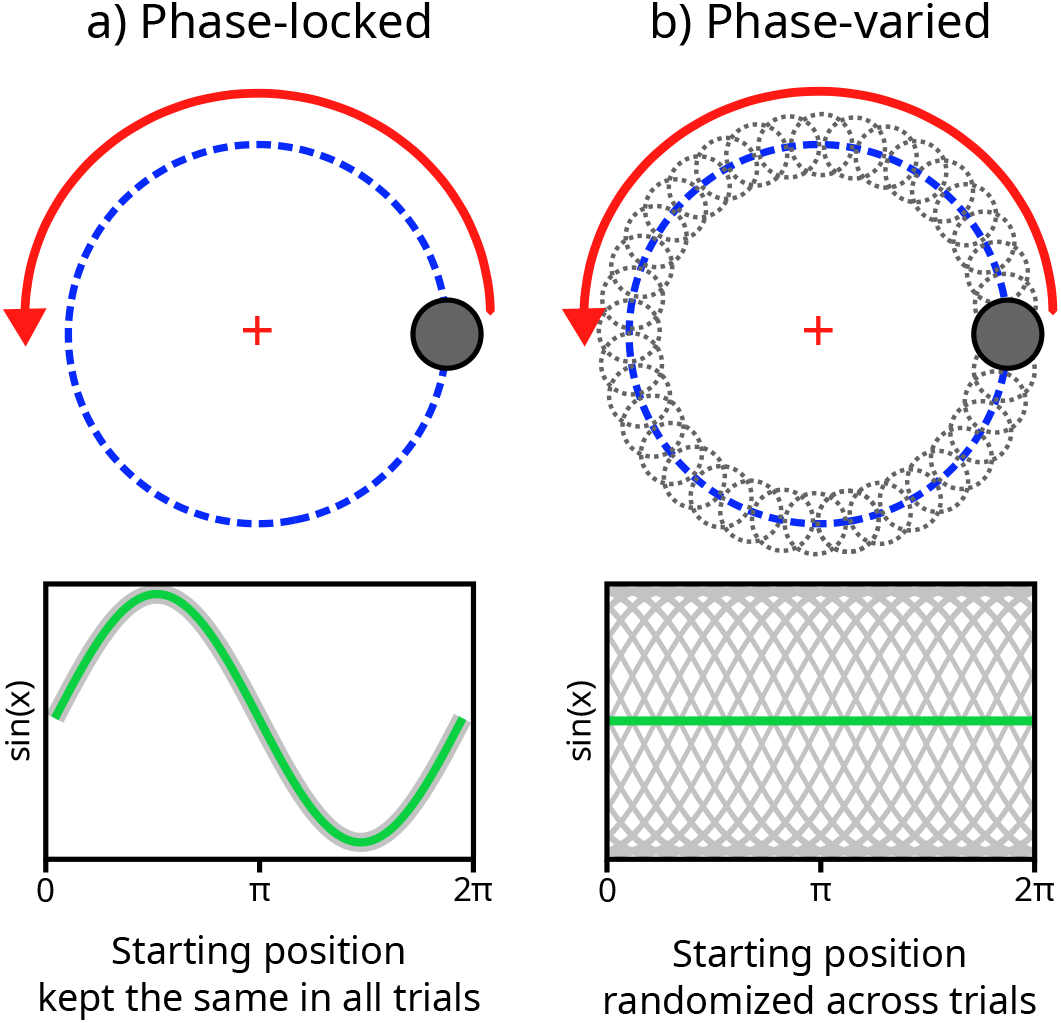
Phase manipulation of the polar angle modulation. **a)** In the phase-locked condition, the dot consistently initiated its rotation from the same polar angle across trials, resulting in a consistent phase for the polar angle modulation. The sine functions (green lines) in the second row are provided for illustrative purposes, demonstrating how the dot’s starting angle influences the phase of the polar angle modulation. Although the actual signal evoked by the polar angle modulation may not be sinusoidal, its phase is expected to align with the phase of this modulation. **b)** In the phase-varied condition, the dot’s starting angle was randomly chosen on each trial from the 36 predefined polar angles. Consequently, the polar angle modulation was not phase-locked across trials in the phase-varied condition, which would result in non-phase-locked signals canceling out each other when averaged in the time domain (the straight green line).

The only difference between the two phase conditions was the locked vs. varied phase of the polar angle modulation. All other stimulus properties, including motion speed, direction-switch rate, rotational frequency, and eccentricity were identical between the two conditions. Moreover, these properties were either not expected to induce periodical signal fluctuations as they were not periodically modulated (e.g., motion speed) or were expected to yield identical frequency components in both conditions as their modulations were phase-locked in all trials (e.g., direction-switch rate, rotational frequency). For example, even when the dot started its rotation from different angles, it repeated its rotation cycle at exactly 1-second intervals (i.e., rotational frequency). Likewise, the dot’s movement direction was shifted by the same amount at each frame regardless of its starting polar angle (i.e., direction-switch rate). Therefore, we expected that any differences between the frequency components extracted from the two phase conditions would primarily stem from the phase manipulation applied to the polar angle modulation.

Since we were interested in EEG activity sensitive to polar angle modulation, we expected rotation direction to be a defining factor in the profile of the modulated EEG signal. Therefore, we also manipulated the rotation direction of the dot. In both phase conditions, the dot rotated either clockwise (CW) or counterclockwise (CCW), counterbalanced within each condition. Hence, we had 36 CW and 36 CCW trials for each phase condition.

#### Procedure

Each trial started with a fixation cross displayed on a gray background for 1 second, followed by an 11-second presentation of the rotating dot stimulus. After each trial, there was a self-paced break in which the participants pressed a keyboard key to proceed with the next trial. In approximately 14% of the trials (12 trials per block), the rotating dot briefly changed its color for 250 milliseconds at a random time (except during the first and last seconds of the trial). Participants were instructed to fixate on a red cross at the center of the screen, and press keys on the keyboard whenever they noticed a change in the dot’s color (F key if the dot turned red, J if it turned green). The trial was terminated upon response or after the 1-second response window, with a feedback (i.e., correct/incorrect) text displayed for 1 second. These trials acted as catch trials to motivate participants to pay attention to the stimulus.

Trials from PL and PV conditions were presented in two separate blocks, the order of which was counterbalanced between participants. Each block consisted of 72 experimental and 12 catch trials, with an equal number of CW and CCW trials, in random order. Participants took a 3-5 minute break between the two blocks. Prior to the experiment, participants completed a short practice block (2-3 minutes) of 12 catch trials to get accustomed to the task.

#### EEG acquisition

EEG was recorded using 64 Ag/AgCl active electrodes placed on the scalp according to the international 10-10 system (ActiCAP, Brain Products GmbH, Gilching, Germany) with the following modifications: The channels TP10 and FT10 were placed above and below the participants’ right eye to be used as vertical electrooculogram (VEOG) channels, and TP9 was used to replace the otherwise absent Iz channel. BrainVision Recorder software (Brain Products GmbH, Gilching, Germany) was used to record the EEG data at a 1000 Hz sampling rate with channel impedances kept below 20 kΩ.

#### EEG preprocessing

Fieldtrip Toolbox (Oostenveld et al., 2010) and custom scripts were used to preprocess the data. A fourth-order Butterworth bandpass filter (0.5 - 100 Hz) was applied to the continuous data. Power line noise was removed by a multi-notch filter at frequencies 50, 100, and 150 Hz with bandwidths of 1, 2, and 3 Hz respectively. The online reference channel (FCz) was re-added to the data and channels were re-referenced to the average of all channels. Blink artifacts were removed by implementing an Independent Component Analysis (ICA) in Fieldtrip, using EEGLAB’s runica algorithm (Delorme & Makeig, 2004). Bad channels were determined by visual inspection and interpolated (0.37% per participant on average) using neighboring channels. Data were then segmented into 11-second trial epochs and downsampled by a quarter to 250 Hz for computational efficiency. A small number of trials were rejected (M = 1.38%, SD = 2.38) due to hardware-related artifacts (peak amplitude > 500 µV). Data from the catch trials were excluded from all analyses.

#### Frequency domain analysis

We expected position-dependent discrepancies in EEG strength to induce periodic fluctuations in the signal by enhancing or attenuating the strength of the population response at different polar angles of the rotating dot stimulus. Therefore, we expected the phases of these periodic fluctuations to align with the phase of our polar angle modulation. Consequently, we anticipated these fluctuations to exhibit phase-locking across the trials of our PL condition, but not across the trials of the PV condition.

To extract the periodic fluctuations induced by the polar angle modulation, we utilized time-domain averaging and averaged the trials from each of our conditions in the time domain (Trial N *≈* 36). Time-domain averaging eliminates noise while keeping phase-locked signals intact (Trimble, 1968). As a consequence, we expected time-domain averaging to preserve the polar angle modulation frequencies in the phase-locked condition, but eliminate the same frequencies in the phase-varied condition through phase-cancellation. As previously discussed (see, Apparatus and stimuli), other stimulus properties were either not expected to induce periodic fluctuations or were expected to induce the same periodic fluctuations in both conditions. Therefore, any frequency components with amplitudes significantly higher in the PL than the PV condition should belong to fluctuations induced by the polar angle modulation alone.

We applied a Fourier transform on the time-averaged data (frequency resolution: 1/11 = 0.09 Hz), and calculated the baseline-subtracted amplitudes by subtracting from the amplitude of each frequency bin the average amplitude of 16 neighboring bins (8 from both sides), with the exclusion of two immediately adjacent bins (one on each side). Since the dot’s polar angle was modulated at 1 Hz, we expected to observe frequency components at 1 Hz and its harmonics. We also compared the overall amplitude of activity distributed across these fundamental and harmonic frequencies. To this end, we summed the baseline-subtracted amplitudes of all harmonic frequencies which were significantly above the noise level (Retter & Rossion, 2016; Retter et al., 2021). Significant harmonics were determined by first grand-averaging the FFT spectra across participants, channels, and conditions, and then calculating Z-scores for each frequency bin with a similar procedure to the baseline-subtracted amplitude calculation (i.e., 16 neighboring bins were used as the baseline for each frequency bin). Only the frequency components up to the sixth harmonic (including the fundamental frequency) passed the threshold of significance (*Z* > 1.64, *p* < 0.05, one-tailed, signal > noise), and thus were included when calculating the summed baseline-subtracted amplitudes for each condition.

We identified two separate ROIs for our analyses. The first included only the Oz channel, which largely covers V1 where the position-dependent discrepancies across different retinotopic areas, such as cortical magnification, are most well-known (Dahlem & Tusch, 2012; Daniel & Whitteridge, 1961; Dow et al., 1981; Gattass et al., 1987; Palmer et al., 2012; Van Essen et al., 1984). The second ROI consisted of 8 posterior channels (Oz, O1, O2, POz, PO3, PO4, PO7, PO8) which roughly encompasses the occipital-parietal areas in which previous studies consistently found frequency components induced by position-modulated stimuli (Aissani et al., 2011; Alp et al., 2017; Nozaradan et al., 2012; Palomares et al., 2012; Pitchaimuthu et al., 2021; Varlet et al., 2023). Separately for each ROI, We ran a 2-by-2 repeated-measures ANOVA (rm-ANOVA) on participants’ summed baseline-subtracted amplitudes with phase (PV vs. PL) and rotation direction (CW vs. CCW) as within-subject factors. For the first ROI, the summed baseline-subtracted amplitudes from the Oz channel were used in the rm-ANOVA. For the second ROI, we averaged the summed baseline-subtracted amplitudes from the 8 channels to quantify and compare the overall response in these posterior channels. All reported p-values for both tests were Bonferroni-corrected for multiple comparisons (i.e., one for each ROI; Keil et al., 2022). Lastly, we calculated the signal-to-noise ratio (SNR = signal/baseline) for all frequency bins using the same baseline as described above, to better illustrate the pattern of harmonic frequencies contained in the evoked signal. SNR is preferred for this purpose as higher frequencies can have a low amplitude (and therefore a low baseline-subtracted amplitude) but still have a high SNR.

## Results

### Behavioral results

The behavioral task in the catch trials served the sole purpose of encouraging participants to pay attention to the stimulus. Therefore, we only required participants to have accuracy above 50% on the catch trials. Overall, participants’ accuracy was high (M = 88.14%, SD = 8.31), and no participant had below-chance accuracy in either of the two experimental blocks (Min. = 58.33%).

### Frequency domain results

Z-scores calculated on the grand-averaged amplitude spectra indicated that harmonics of the polar angle modulation frequency were significant up to the sixth harmonic (see Frequency domain analysis), thereby constituting our frequencies-of-interest for the following analyses. Figure 4 illustrates that these six frequencies are clearly distinguishable from noise in the two PL conditions (CW and CCW) whereas they are at noise levels (SNR ≈ 1) in the PV conditions. A visual inspection of the topographical maps (Figure 5) displaying the summed baseline-subtracted amplitudes for each condition across EEG channels suggests a clear posterior activity in the PL, but not in the PV, conditions.

**Figure 4.**
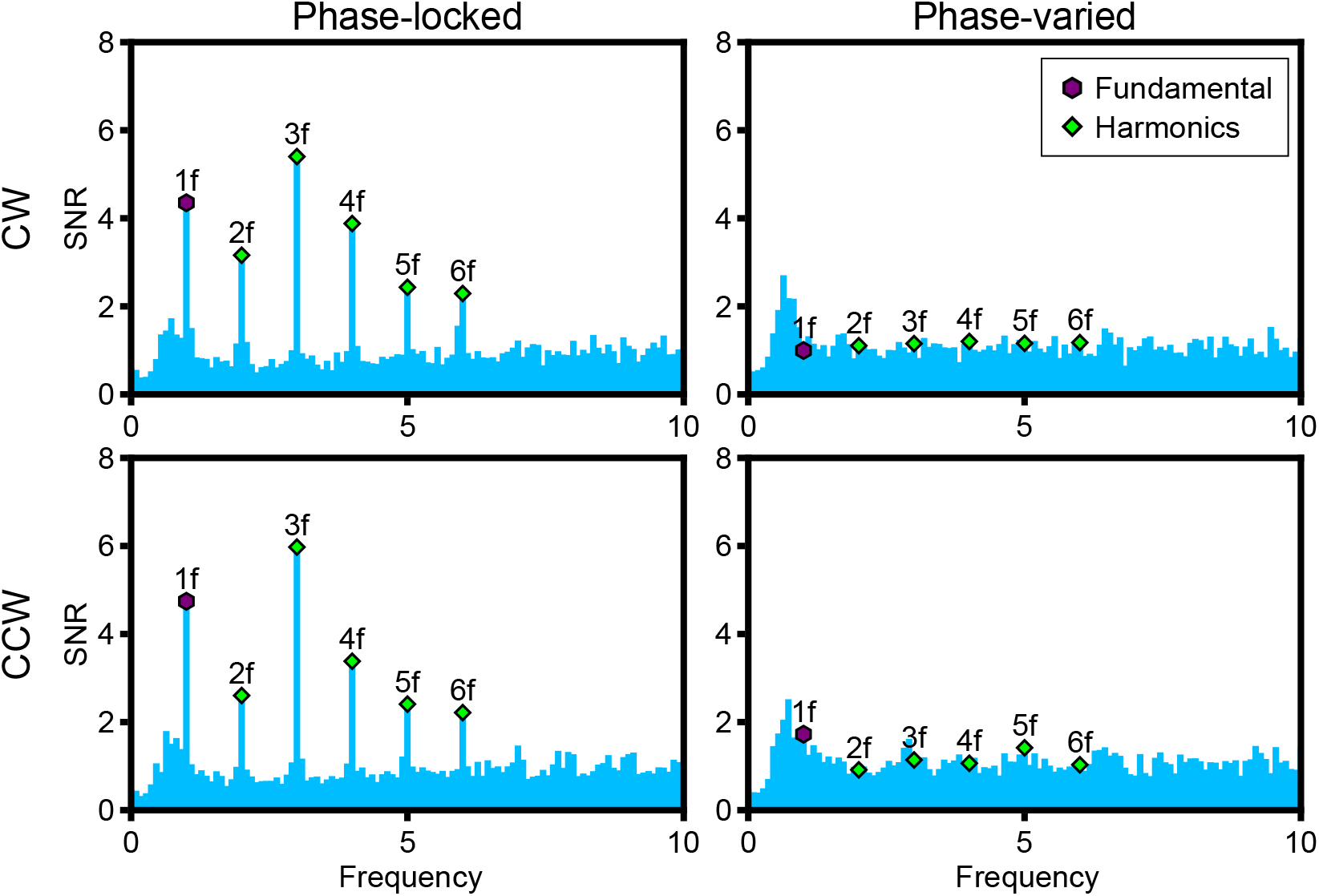
FFT signal-to-noise ratio (SNR) spectra. SNR calculated on the FFT amplitude spectra obtained from time-domain averaged trials for each condition. The figures here reflect the SNR values extracted from the Oz channel, averaged across participants. The first six harmonics of the modulation frequency (1*f*, 2*f*, 3*f*, 4*f*, 5*f*, and 6*f*) exhibit high SNR (> 2) in the phase-locked conditions, but are at noise levels (≈ 1) in the phase-varied conditions.

**Figure 5.**
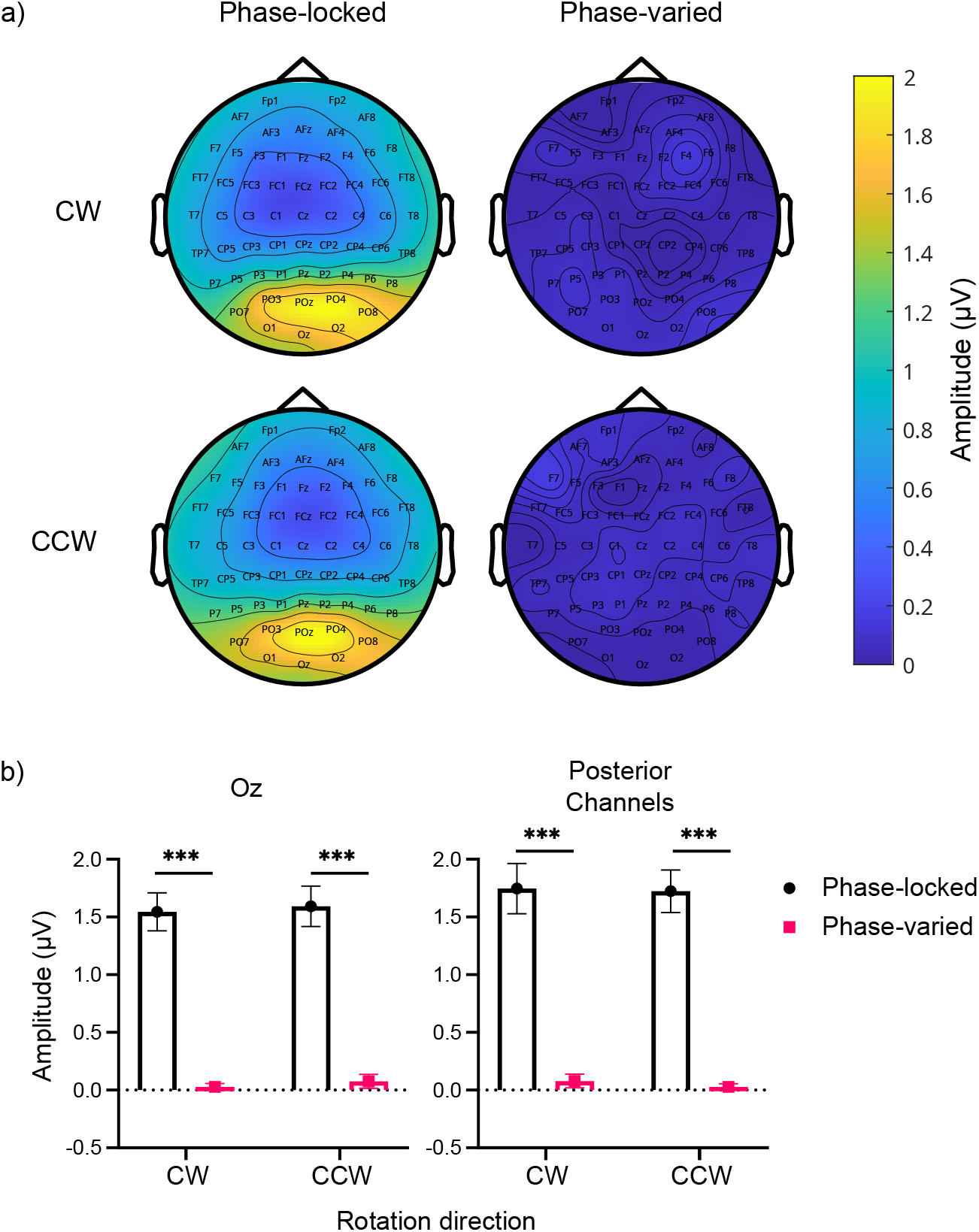
Summed baseline-subtracted amplitudes. **a)** Topographical plots of grand-averaged summed baseline-subtracted amplitudes of the first six harmonics of the modulation frequency (1f, 2f, …, 6f) across all channels. **b)** The summed baseline-subtracted amplitudes of the first six harmonics of the modulation frequency (1f, 2f, …, 6f) are depicted for the two regions-of-interest. Left: Summed baseline-subtracted amplitudes obtained from the Oz channel. Right: Average of the summed baseline-subtracted amplitudes obtained from the eight posterior EEG channels. Error bars indicate the standard error of the mean. Statistical significance is denoted as *** (p < 10^−5^).

A rm-ANOVA on the summed baseline-subtracted amplitudes from the Oz channel (i.e., the first ROI) revealed a significant main effect of phase (*F* (1, 12) = 86.74, *p*_bonf_ < 10^−5^, η^2^_p_ = 0.878). The main effect of rotation direction (*F* (1, 12) = 0.59, *p*_bonf_ = 0.91, η^2^_p_ = 0.047) and the interaction term (*F* (1, 12) = 5.86*10^−5^, *p*_bonf_ = 1, η^2^_p_ = 4.88*10^−6^) were not significant. Figure 5b depicts the distinction among each condition.

Bonferroni-corrected post hoc tests revealed that the summed baseline-subtracted amplitudes from the PL condition were significantly larger than those from the PV condition, within both CW (M_diff_ = 1.516, *p*_bonf_ < 10^−5^) and CCW (M_diff_ = 1.517, *p*_bonf_ < 10^−5^) conditions. CW and CCW conditions did not differ significantly within PV (M_diff_ = 0.048, *p*_bonf_ = 1) or PL (M_diff_ = 0.047, *p*_bonf_ = 1) conditions.

A separate rm-ANOVA on the summed baseline-subtracted amplitudes averaged from the eight posterior channels (i.e., the second ROI) also revealed similar results: The main effect of phase (*F* (1, 12) = 75.77, *p*_bonf_ < 10^−5^, η^2^_p_ = 0.863) was significant, but the main effect of rotation direction (*F* (1, 12) = 0.48, *p*_bonf_ = 1, η^2^_p_ = 0.039) and the interaction term (*F* (1, 12) = 0.064, *p*_bonf_ = 1, η^2^_p_ = 0.005) were not. The PL condition had larger summed baseline-subtracted amplitudes than the PV condition in both CW (*M*_*diff*_ = 1.67, *p*_bonf_ < 10^−5^) and CCW (*M*_*diff*_ = 1.69, *p*_bonf_ < 10^−5^) conditions (see Figure 5b). There was no significant difference between the CW and CCW conditions within PV (*M*_*diff*_ = 0.05, *p*_bonf_ = 1) or PL (*M*_*diff*_ = 0.023, *p*_bonf_ = 1) conditions.

#### Experiment 2: Eccentricity modulation

Results from Experiment 1 revealed frequency components that are contingent on the phase of the polar angle modulation, and therefore originate from a position-dependent modulation of the EEG signal. These results are in line with our hypothesis that systematic position-dependent discrepancies in the EEG strength can yield frequency components by interacting with a position-modulated stimulus. In Experiment 2, we investigated whether a modulation of stimulus eccentricity can drive similar results.

## Materials and Methods

Materials and methods were identical to Experiment 1, with the exceptions outlined below.

### Participants

13 Sabanci University students (8 female, 5 male; mean age = 21.38 years, SD = 0.5) with normal or corrected-to-normal vision were recruited in exchange for course credit. The sample size was based on previous studies as in Experiment 1. A sensitivity analysis revealed that this sample size was sufficiently powered (power = .80) to detect effect sizes larger than 0.73 (Cohen’s d, one-tailed paired-samples t-test; see, Frequency domain analysis).

### Apparatus and Stimuli

In Experiment 2, we employed the same rotating dot stimulus but adjusted the position of the rotation trajectory by shifting it in eight different directions (shift direction angles: 0°to 315°in increments of 45°). As a consequence, the center of the rotation trajectory was always positioned 2 dva away from the fixation cross, which allowed the dot’s eccentricity to be modulated between 3 and 7 dva during its rotation. The phase of this eccentricity modulation depended on the position from which the dot started its rotation. More specifically, the phase of the eccentricity modulation was determined by the relative angle between the dot’s initial position and the axis of the trajectory shift (Figure 6). Using this property, we kept the phase of the eccentricity modulation constant across trials in the phase-locked (PL) eccentricity modulation condition and varied it in the phase-varied (PV) condition (range: 0°to 315°in steps of 45°). The constant phase used for the eccentricity modulation in the PL condition was randomized between participants by randomly assigning each participant a predetermined relative starting angle. This meant that the relative starting angle of the dot in the PL condition, and therefore the phase of the eccentricity modulation, was different for each participant (but still constant across the PL trials within participants).

**Figure 6.**
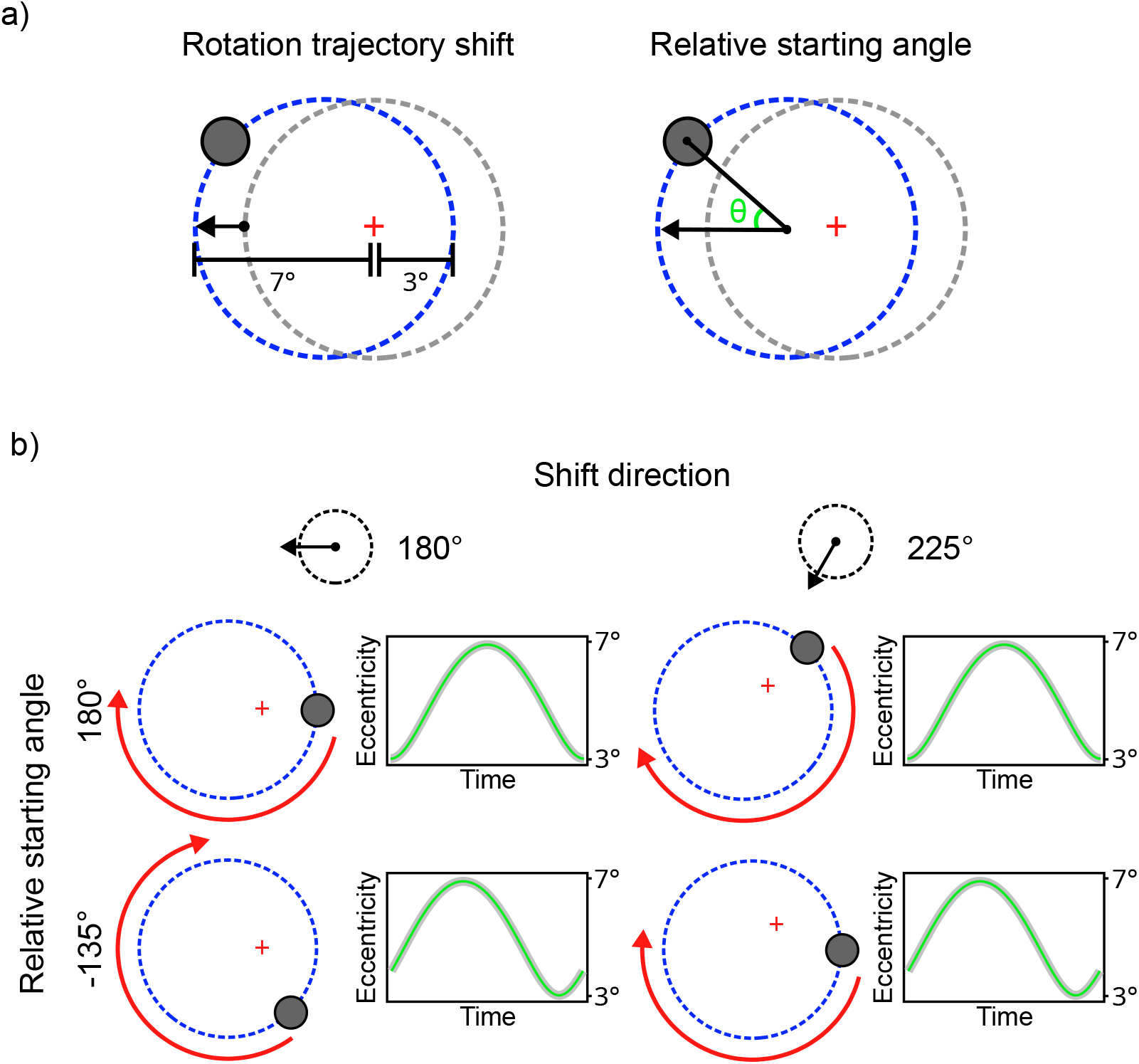
Rotation trajectory shift and the phase of the eccentricity modulation. **a)** Left: The circular rotation trajectory was shifted by 2 dva. As a result, the dot’s eccentricity changed from 3 to 7 dva during its rotation. Right: Relative starting angle of the dot defined as the angle between the vector of the rotation trajectory shift and the dot’s initial position (shift angle - dot’s polar angle). This relative starting angle determined the phase of the eccentricity modulation. **b)** The phase of the eccentricity modulation was determined by the relative starting angle of the dot, even when different shift angles were applied to the rotation trajectories. The left and right columns represent conditions where the rotation trajectories were shifted in different directions, yet the phases of the eccentricity modulation remained the same. This consistency was achieved by maintaining the same relative starting angle across different shift directions.

Shifting the rotation trajectory in eight different directions allowed for varying the phase of the polar angle modulation in both phase conditions, while still enabling a phase-locked eccentricity modulation in the PL condition. Therefore, we expected to observe only the frequency components introduced by the eccentricity modulation, as those introduced by the polar angle modulation would be eliminated through phase-cancelation. As a consequence of the cancellation of polar angle-dependent signals, any influence of the dot’s rotation direction (CW and CCW) was also expected to be eliminated. Therefore, to increase the number of trials for our phase conditions, we omitted the rotation direction conditions in Experiment 2. Instead, we maintained a constant rotation direction (either clockwise or counterclockwise) across all trials, with the specific direction randomly predetermined for each participant.

Each trial lasted 10 seconds and contained 10 full rotations of the dot. Phase-locked and phase-varied conditions were presented in two different blocks similar to Experiment 1. Each block had 76 trials, 12 of which were catch trials. The task in the catch trials was identical to Experiment 1.

### Frequency domain analysis

Bad channels were interpolated (0.5% per subject on average) using neighboring channels, and a small number of trials (M = 0.96%, SD = 1.15) with hardware-related artifacts were excluded from the analyses. Later, trials from the PL and PV conditions were averaged separately in the time domain (Trial N *≈* 64). Following the same Z-score calculation procedure and significance criterion as in Experiment 1 (baseline: 14 neighboring bins for each frequency), only the fundamental frequency component (1 Hz) was found significant. Therefore, we used the baseline-subtracted amplitudes of the fundamental frequency to compare the two conditions in two paired-samples t-tests for the two ROIs. Since we expected the PL condition to produce large baseline-subtracted amplitudes and the PV condition around the noise level (baseline-subtracted amplitude *≈* 0), we ran one-tailed tests.

## Results

### Behavioral results

Participants’ accuracy on the behavioral task was similar to Experiment 1 (M = 84.62%, SD = 9.22, Min. = 58.33%).

### Frequency domain results

Z-scores calculated on the grand-averaged amplitude spectra revealed that only the fundamental frequency was significantly larger than the noise (*Z* = 2.75, *p* = .003, one-tailed, signal > noise). Figure 7 shows that the fundamental frequency had a large SNR in the PL condition, but was at noise levels in the PV condition (SNR *≈* 1).

**Figure 7.**
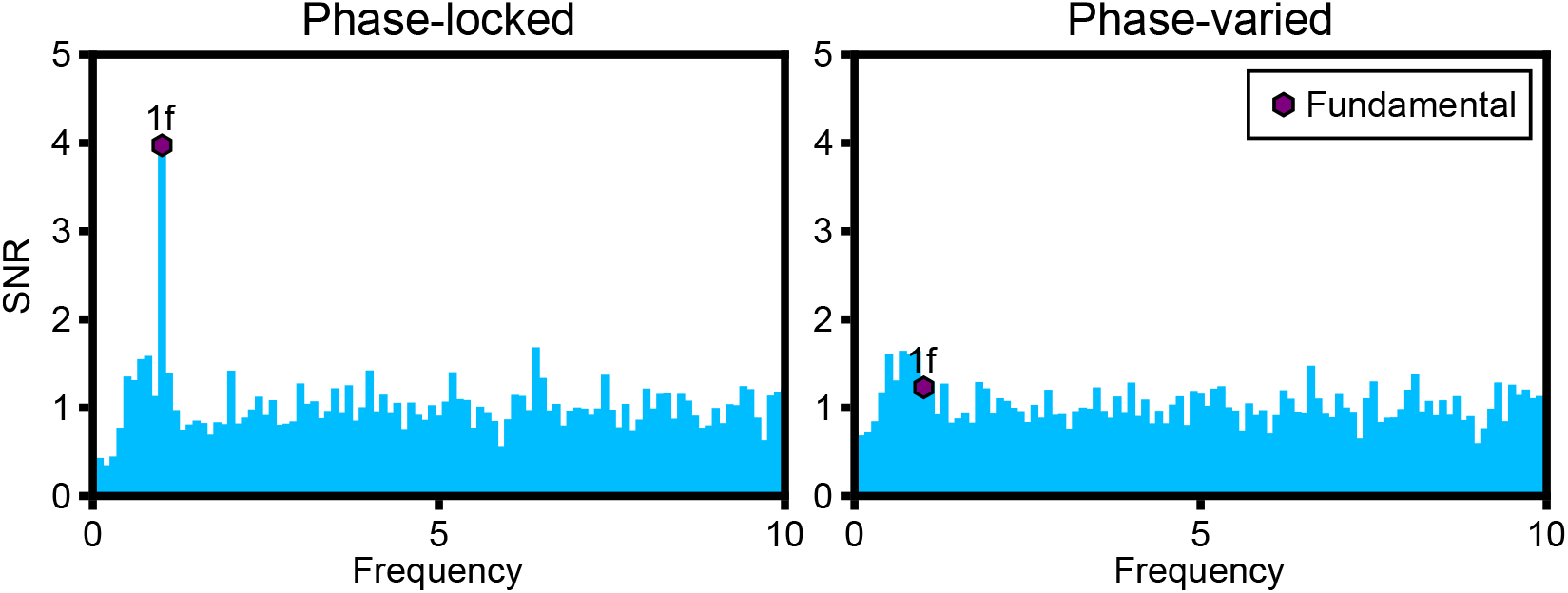
FFT signal-to-noise ratio (SNR) spectra. SNR calculated on the FFT amplitude spectra obtained from time-domain averaged trials for each condition, from the Oz channel. The fundamental modulation frequency (1f) exhibits a high SNR in the phase-locked condition, whereas its SNR is at noise levels in the phase-varied condition.

Topographical maps of the baseline-subtracted amplitudes (Figure 8b) suggest that phase-locked eccentricity modulation elicited activity in the posterior channels, whereas no particular activity was observed in the PV condition.

**Figure 8.**
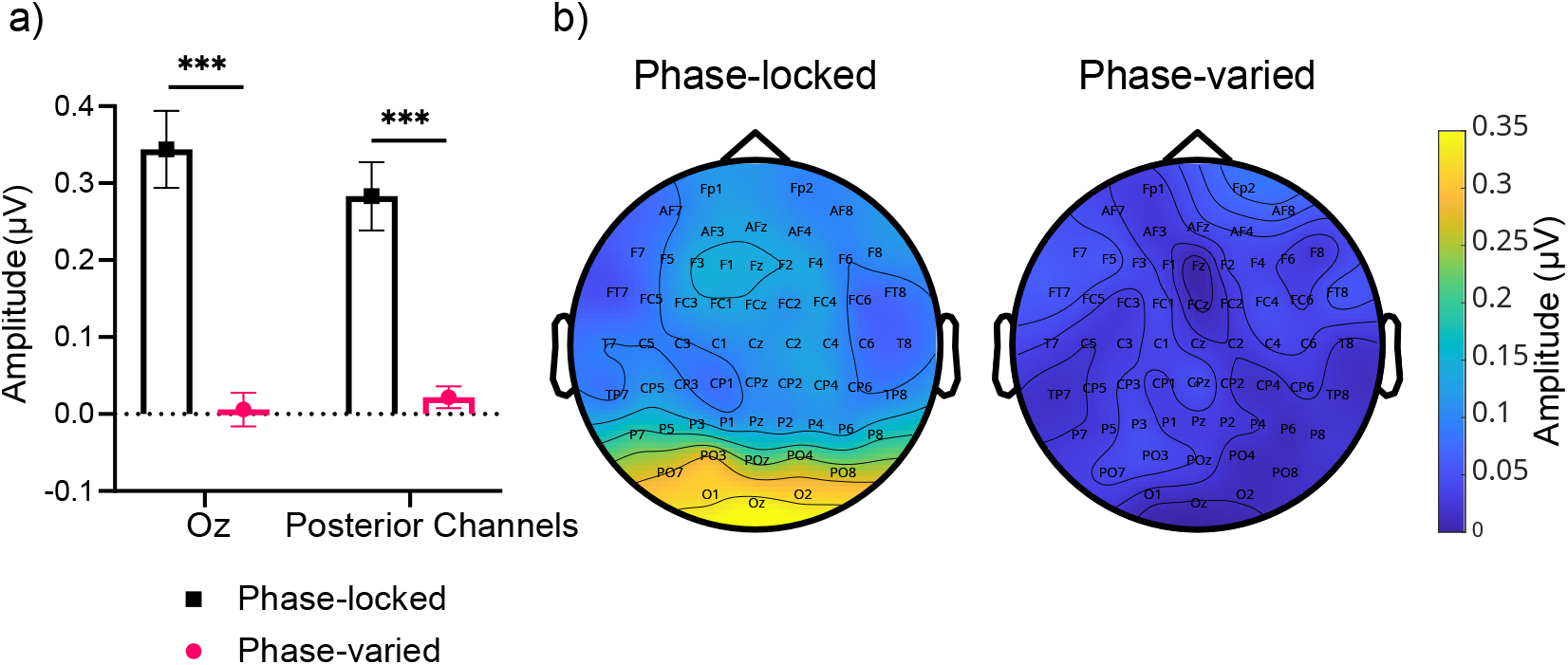
Baseline-subtracted amplitudes. **a)** The baseline-subtracted amplitudes of the modulation frequency (1*f*) are depicted for the two regions-of-interest: Baseline-subtracted amplitudes obtained from the Oz channel, and average of the baseline-subtracted amplitudes obtained from the eight posterior EEG channels. Error bars indicate the standard error of the mean. Statistical significance is denoted as *** (*p* < .001). **b)** Topographical maps of grand-averaged baseline-subtracted amplitudes of the fundamental modulation frequency (1*f*) across all channels.

We ran two separate one-tailed (i.e., PL > PV) paired-samples t-tests on the two ROIs. The fundamental frequency had a significantly larger baseline-subtracted amplitude in the PL condition for both the Oz channel (*t*(12) = 5.17, *p*_bonf_ < .001, *d*= 1.43), and the average of the eight posterior channels (*t*(12) = 5.12, *p*_bonf_ < .001, *d*= 1.42), compared to the PV condition (Figure 8a).

## Discussion

We hypothesized that position-dependent factors that influence the strength of signals measured from different retinotopic populations, such as their volume and depth, could introduce consistent fluctuations in EEG signals under certain circumstances. We investigated whether the signal fluctuations driven purely by such position-dependent factors could yield frequency components when no other stimulus property was periodically modulated. Our simulations revealed that position-dependent factors could interact with a position-modulated stimulus, generating signal fluctuations that produce distinct frequency components. These signal fluctuations are strictly phase-coupled to the stimulus’s position modulation. Furthermore, the frequency components they produce depend on both the trajectory of the position modulation and the way position-dependent factors influence signals measured from different retinotopic populations (i.e., Retinotopic Weight Function).

In two subsequent experiments, we modulated the position of a singleton shape by separately modulating its polar angle (Experiment 1) and eccentricity (Experiment 2). For both types of position modulation, we had a phase-locked condition in which the phase of the modulation was synchronized across trials, and a phase-varied condition where the phase of the modulation was randomized across trials. Moreover, we kept all other stimulus properties constant between the two conditions so that these properties would either yield no frequency components, or would yield identical frequency components in both conditions. Our goal was to selectively eliminate signal fluctuations produced by position-dependent factors through phase-cancellation in the phase-varied conditions. This way, the frequency components that differ between the phase-locked and phase-varied conditions would indicate the existence of position-dependent factors that can produce consistent periodic signal fluctuations by interacting with a position-modulated stimulus.

In line with our hypothesis, we observed clear frequency components at the position modulation frequencies in both experiments when the phases of the polar angle and eccentricity modulations were synchronized across trials. However, when the phases of these modulations were randomized, the same frequency components were indistinguishable from noise.

Frequency components in SSVEP studies typically exhibit narrowband characteristics, as they arise from periodic fluctuations in neural activity in response to stimulus properties modulated at specific frequencies. In our study, the only stimulus property that was periodically modulated was its position. Consequently, the observed frequency components are best explained by variations in the structural properties of different retinotopic populations captured in EEG signals, rather than by the activity of feature-tuned neurons.

This position-dependent modulation of the EEG signal can be readily explained by the retinotopic discrepancies in cortical properties that influence the strength of the measured EEG signals (e.g., cortical volume: Schaul, 1998; cortical depth: Butler et al., 2019; cancellation between simultaneously active sources: Ahlfors et al., 2010). Simple amplification or attenuation of signals across different retinotopic areas—for instance, due to variations in the size or depth of the activated regions—can produce signal fluctuations based on stimulus position. Therefore, any set of structural factors that systematically influences EEG amplitude could potentially generate distinguishable frequency components by introducing position-dependent signal fluctuations.

For instance, due to the non-uniform mapping of the visual field to the visual cortex, parts of the visual field are processed by disproportionately larger brain areas, a property known as cortical magnification (Daniel & Whitteridge, 1961; Horton & Hoyt, 1991; Schwartz, 1980). In our study, the position-modulated stimulus may have activated brain areas of varying sizes depending on the stimulus position, leading to differences in EEG amplitudes as a result of cortical magnification. In this case, the observed frequency components would reflect the differences in the size of the activated brain regions and the total number of neurons within them, rather than fluctuations in the responses of neurons tuned to the stimulus.

The key point here is not that any one structural factor solely explains these results, but that such structural factors—either collectively or independently—can generate frequency components unrelated to the functional sources of EEG fluctuations. This finding challenges a core assumption of SSVEP methodology, which posits that the frequency components observed in response to a periodically modulated stimulus are a direct reflection of evoked neuronal activity processing the specific stimulus property that is modulated (Norcia et al., 2015). In contrast, the frequency components we report do not arise from any specific neural processing mechanism. Instead, structural factors superimpose on signals originating from intrinsic neural processing and modify those signals as an independent source of amplitude modulation. As a result, while these frequency components may resemble typical SSVEP components, they do not provide insights into the brain’s processing of visual information. Rather, they emerge as a byproduct of the brain’s structural organization.

The frequency components generated by position-dependent factors present a particular challenge when they closely resemble those expected to result from specific neural mechanisms or feature-tuned populations. For instance, in SSVEP studies focusing on visual motion processing, researchers often employ stimuli composed of moving visual elements whose positions undergo repetitive modulation along a fixed path of motion (e.g., Aissani et al., 2011; Nozaradan et al., 2012; Pitchaimuthu et al., 2021; Varlet et al., 2023). Some of these studies have argued that the frequency components elicited in response to such stimuli reflect the modulation in the activity of motion-sensitive neural populations, including those sensitive to motion direction (e.g., Aissani et al., 2011; although their results based on intermodulation components cannot be readily explained by structural factors) or repetitive motion patterns (e.g., Aissani et al., 2011; Pitchaimuthu et al., 2021). However, with such motion stimuli, the motion-related characteristics are inherently linked to position modulation. Consequently, position-dependent periodic fluctuations could lead to similar frequency components either independently or in conjunction with the signal fluctuations introduced by the modulated activity of motion-sensitive populations. This complicates the interpretation of results from these studies, making it challenging to attribute the observed frequency components solely to specific neural mechanisms or populations.

Interestingly, our results do not implicate specific neural responses sensitive to repeating motion patterns, or “visual beats”, that previous studies have reported (Nozaradan et al., 2012; Pitchaimuthu et al., 2021). In our experiments, the singleton dot stimulus always completed a full cycle every 1 second in both phase-locked and phase-varied conditions. If there were neural responses specific to the periodicity of motion, we would have observed them in both of our phase conditions. Instead, we only observed significant frequency components when the modulations of stimulus position were phase-locked across trials. This suggests that these frequency components were not driven by neural mechanisms sensitive to the periodicities in motion, but rather by position-dependent modulation of the EEG signal.

Another interesting aspect of our data is the differing patterns of frequency components resulting from our polar angle and eccentricity modulations. The polar angle modulation in Experiment 1 generated complex periodic waveforms that included the first six harmonics of the modulation frequency (1*f*, 2*f*, 3*f*, 4*f*, 5*f*, and 6*f*). In contrast, the eccentricity modulation in Experiment 2 produced simpler, sinusoidal waveforms, containing only the modulation frequency (1*f*). This difference is intriguing because it likely reflects how the position-dependent factors influencing EEG strength vary as a function of polar angle compared to eccentricity.

The frequency components we observed in our simulations always depended on an interaction between the type of the position modulation and the Retinotopic Weight Function. As such, different types of position modulation led to different frequency components even for the same Retinotopic Weight Function. In the experimental data, the actual Retinotopic Weight Function of the visual system is assumed to be consistent across the two experiments. Therefore, the difference in the observed frequency components must be due to the different types of position modulation.

In Experiment 1, the stimulus rotated steadily along an imaginary circle, allowing variations in the Retinotopic Weight Function across polar angles to be directly captured in the signal. A constant population response across polar angles would yield a flat signal with no frequency components, while variations across polar angles would result in observable frequency components that reflect the specific pattern of the variation. In Experiment 2, both polar angle and eccentricity were modulated, but only the eccentricity modulation was phase-locked across trials, which followed a sinusoidal pattern. Thus, if the Retinotopic Weight Function varied linearly with eccentricity, the resulting signal would reflect the sinusoidal pattern of the eccentricity modulation. In contrast, a non-linear variation across eccentricity would produce a more complex, non-sinusoidal signal containing harmonics of the modulation frequency.

In this regard, our results suggest that the actual Retinotopic Weight Function of the visual system is one that varies linearly across eccentricity—at least within the range of our eccentricity modulation (3-7 dva)—and non-linearly across polar angles. This observation aligns with previous findings indicating that cortical magnification factors and population receptive field sizes decrease roughly linearly within this eccentricity range (i.e., 3-7 dva; Daniel and Whitteridge, 1961; Dumoulin and Wandell, 2008) while they vary non-linearly with polar angle (Silva et al., 2018). However, it remains to be elucidated how other structural factors, aside from cortical magnification, may contribute to the differences in results between the two experiments.

We have proposed that the frequency components observed in our results reflect the brain’s structure rather than its function. However, could these position-dependent frequencies arise from functional rather than structural sources? Specifically, might these frequency components be driven by neural mechanisms that produce stronger or weaker responses based on stimulus position?

The observed frequency components were most prominent in posterior visual areas, where neurons possess small, highly overlapping receptive fields. For example, a neuron in the striate cortex typically has a receptive field spanning less than 1 degree of visual angle (Dow et al., 1981; Schiller et al., 1976). With our position-modulated stimulus, the entire receptive field of such a neuron would be traversed in less than ∼30 milliseconds. This suggests that the full range of response variations for this neuron would occur within a time window far shorter than the periods of the frequency components in our data (∼160–1000 ms). Furthermore, the overlapping receptive fields of neighboring neurons would generate out-of-phase responses, effectively canceling out these functional variations as the degree of overlap increases. Therefore, large-scale position-dependent signal variations observed in our results cannot originate from fluctuations in individual neuron responses in these early visual areas.

An alternative explanation is that these position-dependent signal variations are mediated by feedback from higher visual areas, where neurons have larger receptive fields. For instance, object-selective neurons in the monkey IT cortex (analogous to the LOC in humans) have receptive fields spanning ∼50 dva and exhibit enhanced responses to foveal stimuli (Rolls et al., 2003). Feedback from such neurons to lower-level visual areas could partially explain our findings, particularly regarding eccentricity modulation. However, it remains unclear whether the foveal bias observed in IT neurons originates independently or arises from increased feedforward input from lower-level areas driven by the aforementioned structural factors such as cortical magnification. Consequently, it is still uncertain whether position-dependent responses can emerge in high-level visual areas independently of structural influences.

One of the key advantages of SSVEPs is the ability to isolate and track the activity of specific neural populations tuned to specific properties of sensory inputs. However, achieving this requires careful consideration of the targeted neural populations that will be influenced by experimental manipulations, as well as the structural and physical factors governing the translation of neural activity into EEG signals. Our study highlights a scenario in which the observed frequency components do not reflect the tuned activity of a selective neural population of interest, but instead capture the inherent discrepancies in EEG strength that vary systematically across different retinotopic areas due to position-dependent factors. If not adequately addressed in the experimental design, these discrepancies could introduce confounding influences into the data, and lead to the misinterpretation of observed frequency components, particularly in the context of position-modulated stimuli. Furthermore, they could also cause variations in response amplitudes across different stimuli if those stimuli are consistently positioned at different positions on the retina.

To mitigate these issues, we propose that researchers should give due consideration to these position-dependent factors when designing SSVEP studies. It is crucial to ensure that the stimuli being compared do not have consistent differences in retinotopic position. Furthermore, in experiments involving moving stimuli, it is essential to clearly distinguish between frequency components arising from the visual processes of interest from those generated by position-dependent factors. In our phase-varied conditions, we randomized the phase of our position modulations while keeping the modulations in other motion-related stimulus properties synchronized across trials. This enabled us to eliminate the frequency components arising from position modulation through phase-cancellation. Future studies can employ similar approaches to effectively isolate the main activity of interest from the influence of position-dependent effects.

## Acknowledgements

This study was funded by the TUBITAK 3501 Career Development Grant (220K038) awarded to NA by the Scientific and Technological Research Institution of Turkey.

## Ethics approval

All experimental procedures were reviewed and approved by the Sabanci University Research Ethics Council.

## Consent to participate

All participants signed informed consent prior to the experiment session.

## Consent for publication

All participants signed informed consent for their anonymized data to be published.

## Availability of data and materials

The experiment was not preregistered. All data and materials can be found at the following OSF repository: https://osf.io/mqb49/?view_only=43e71e2549d8485baa73f1ebc82f230a

## References

Ahlfors, S. P., Han, J., Lin, F.-H., Witzel, T., Belliveau, J. W., Hämäläinen, M. S., & Halgren, E. (2010). Cancellation of EEG and MEG Signals Generated by Extended and Distributed Sources. Human Brain Mapping, 31 (1), 140–149. 10.1002/hbm.20851

Aissani, C., Cottereau, B., Dumas, G., Paradis, A.-L., & Lorenceau, J. (2011). Magnetoencephalographic Signatures of Visual Form and Motion Binding. Brain Research, 1408, 27–40. 10.1016/j.brainres.2011.05.051

Allen, D., Tyler, C. W., & Norcia, A. M. (1996). Development of Grating Acuity and Contrast Sensitivity in the Central and Peripheral Visual Field of the Human Infant. Vision Research, 36 (13), 1945–1953. 10.1016/0042-6989(95)00257-x

Allman, J. M., & Kaas, J. H. (1971). Representation of the Visual Field in Striate and Adjoining Cortex of the Owl Monkey (Aotus Trivirgatus). Brain Research, 35 (1), 89–106. 10.1016/0006-8993(71)90596-8

Alp, N., Kogo, N., Van Belle, G., Wagemans, J., & Rossion, B. (2016). Frequency tagging yields an objective neural signature of Gestalt formation. Brain and Cognition, 104, 15–24. 10.1016/j.bandc.2016.01.008

Alp, N., Kohler, P. J., Kogo, N., Wagemans, J., & Norcia, A. M. (2018). Measuring Integration Processes in Visual Symmetry with Frequency-Tagged EEG. Scientific Reports, 8 (1), 6969. 10.1038/s41598-018-24513-w

Alp, N., Nikolaev, A. R., Wagemans, J., & Kogo, N. (2017). EEG Frequency Tagging Dissociates between Neural Processing of Motion Synchrony and Human Quality of Multiple Point-Light Dancers. Scientific Reports, 7 (1), 44012. 10.1038/srep44012

Alp, N., & Ozkan, H. (2022). Neural Correlates of Integration Processes during Dynamic Face Perception. Scientific Reports, 12 (1), 118. 10.1038/s41598-021-02808-9

Butler, R., Bernier, P.-M., Mierzwinski, G. W., Descoteaux, M., Gilbert, G., & Whittingstall, K. (2019). Cortical Distance, Not Cancellation, Dominates Inter-Subject EEG Gamma Rhythm Amplitude. NeuroImage, 192, 156–165. 10.1016/j.neuroimage.2019.03.010

Dahlem, A. M., & Tusch, J. (2012). Predicted Selective Increase of Cortical Magnification Due to Cortical Folding. The Journal of Mathematical Neuroscience, 2 (1), 14. 10.1186/2190-8567-2-14

Daniel, P. M., & Whitteridge, D. (1961). The Representation of the Visual Field on the Cerebral Cortex in Monkeys. The Journal of Physiology, 159 (2), 203–221.

Delorme, A., & Makeig, S. (2004). EEGLAB: An Open Source Toolbox for Analysis of Single-Trial EEG Dynamics Including Independent Component Analysis. Journal of Neuroscience Methods, 134 (1), 9–21. 10.1016/j.jneumeth.2003.10.009

Dow, B. M., Snyder, A. Z., Vautin, R. G., & Bauer, R. (1981). Magnification Factor and Receptive Field Size in Foveal Striate Cortex of the Monkey. Experimental Brain Research, 44 (2), 213–228. 10.1007/BF00237343

Dumoulin, S. O., & Wandell, B. A. (2008). Population receptive field estimates in human visual cortex. NeuroImage, 39 (2), 647–660. 10.1016/j.neuroimage.2007.09.034

Gattass, R., Sousa, A. P. B., & Rosa, M. G. P. (1987). Visual Topography of V1 in the Cebus Monkey. Journal of Comparative Neurology, 259 (4), 529–548. 10.1002/cne.902590404

Gundlach, C., & Müller, M. M. (2013). Perception of illusory contours forms intermodulation responses of steady state visual evoked potentials as a neural signature of spatial integration. Biological Psychology, 94 (1), 55–60. 10.1016/j.biopsycho.2013.04.014

Horton, J. C., & Hoyt, W. F. (1991). The Representation of the Visual Field in Human Striate Cortex. A Revision of the Classic Holmes Map. Archives of Ophthalmology (Chicago, Ill.: 1960), 109 (6), 816–824. 10.1001/archopht.1991.01080060080030

Keil, A., Bernat, E. M., Cohen, M. X., Ding, M., Fabiani, M., Gratton, G., Kappenman, E. S., Maris, E., Mathewson, K. E., Ward, R. T., & Weisz, N. (2022). Recommendations and Publication Guidelines for Studies Using Frequency Domain and Time-Frequency Domain Analyses of Neural Time Series. Psychophysiology, 59 (5), e14052. 10.1111/psyp.14052

Liu, S., Zhang, D., Liu, Z., Liu, M., Ming, Z., Liu, T., Suo, D., Funahashi, S., & Yan, T. (2022). Review of brain–computer interface based on steady-state visual evoked potential. Brain Science Advances, 8 (4), 258–275. 10.26599/BSA.2022.9050022

Norcia, A. M., Appelbaum, L. G., Ales, J. M., Cottereaur, B. R., & Rossion, B. (2015). The Steady State VEP in Research. Journal of Vision, 15 (6), 1–46. 10.1167/15.6.4.doi

Nozaradan, S., Peretz, I., & Mouraux, A. (2012). Steady-State Evoked Potentials as an Index of Multisensory Temporal Binding. NeuroImage, 60 (1), 21–28. 10.1016/j.neuroimage.2011.11.065

Oostenveld, R., Fries, P., Maris, E., & Schoffelen, J.-M. (2010). FieldTrip: Open Source Software for Advanced Analysis of MEG, EEG, and Invasive Electrophysiological Data. Computational Intelligence and Neuroscience, 2011, e156869. 10.1155/2011/156869

Palmer, C. R., Chen, Y., & Seidemann, E. (2012). Uniform Spatial Spread of Population Activity in Primate Parafoveal V1. Journal of Neurophysiology, 107 (7), 1857–1867. 10.1152/jn.00117.2011

Palomares, M., Ales, J. M., Wade, A. R., Cottereau, B. R., & Norcia, A. M. (2012). Distinct Effects of Attention on the Neural Responses to Form and Motion Processing: A SSVEP Source-Imaging Study. Journal of Vision, 12 (10), 15. 10.1167/12.10.15

Peirce, J., Gray, J. R., Simpson, S., MacAskill, M., Höchenberger, R., Sogo, H., Kastman, E., & Lindeløv, J. K. (2019). PsychoPy2: Experiments in Behavior Made Easy. Behavior Research Methods, 51 (1), 195–203. 10.3758/s13428-018-01193-y

Pitchaimuthu, K., Dormal, G., Sourav, S., Shareef, I., Rajendran, S. S., Ossandón, J. P., Kekunnaya, R., & Röder, B. (2021). Steady State Evoked Potentials Indicate Changes in Nonlinear Neural Mechanisms of Vision in Sight Recovery Individuals. Cortex; a Journal Devoted to the Study of the Nervous System and Behavior, 144, 15–28. 10.1016/j.cortex.2021.08.001

Regan, D. (1977). Steady-State Evoked Potentials. Journal of the Optical Society of America, 67 (11), 1475–1489. 10.1364/josa.67.001475

Retter, T. L., & Rossion, B. (2016). Uncovering the Neural Magnitude and Spatio-Temporal Dynamics of Natural Image Categorization in a Fast Visual Stream. Neuropsychologia, 91, 9–28. 10.1016/j.neuropsychologia.2016.07.028

Retter, T. L., Rossion, B., & Schiltz, C. (2021). Harmonic Amplitude Summation for Frequency-tagging Analysis. Journal of Cognitive Neuroscience, 33 (11), 2372–2393. 10.1162/jocn_a_01763

Rice, J. K., Rorden, C., Little, J. S., & Parra, L. C. (2013). Subject Position Affects EEG Magnitudes. NeuroImage, 64, 476–484. 10.1016/j.neuroimage.2012.09.041

Rolls, E. T., Aggelopoulos, N. C., & Zheng, F. (2003). The Receptive Fields of Inferior Temporal Cortex Neurons in Natural Scenes. The Journal of Neuroscience, 23 (1), 339–348. 10.1523/JNEUROSCI.23-01-00339.2003

Rossion, B., Torfs, K., Jacques, C., & Liu-Shuang, J. (2015). Fast Periodic Presentation of Natural Images Reveals a Robust Face-Selective Electrophysiological Response in the Human Brain. Journal of Vision, 15 (1), 18. 10.1167/15.1.18

Sartucci, F., Borghetti, D., Bocci, T., Murri, L., Orsini, P., Porciatti, V., Origlia, N., & Domenici, L. (2010). Dysfunction of the magnocellular stream in Alzheimer’s disease evaluated by pattern electroretinograms and visual evoked potentials. Brain Research Bulletin, 82 (3), 169–176. 10.1016/j.brainresbull.2010.04.001

Schaul, N. (1998). The Fundamental Neural Mechanisms of Electroencephalography. Electroencephalography and Clinical Neurophysiology, 106 (2), 101–107. 10.1016/S0013-4694(97)00111-9

Schiller, P. H., Finlay, B. L., & Volman, S. F. (1976). Quantitative studies of single-cell properties in monkey striate cortex. I. Spatiotemporal organization of receptive fields. Journal of Neurophysiology, 39 (6), 1288–1319. 10.1152/jn.1976.39.6.1288

Schwartz, E. L. (1980). Computational Anatomy and Functional Architecture of Striate Cortex: A Spatial Mapping Approach to Perceptual Coding. Vision Research, 20 (8), 645–669. 10.1016/0042-6989(80)90090-5

Silberstein, R. B., Line, P., Pipingas, A., Copolov, D., & Harris, P. (2000). Steady-state visually evoked potential topography during the continuous performance task in normal controls and schizophrenia. Clinical Neurophysiology, 111 (5), 850–857. 10.1016/S1388-2457(99)00324-7

Silva, M. F., Brascamp, J. W., Ferreira, S., Castelo-Branco, M., Dumoulin, S. O., & Harvey, B. M. (2018). Radial Asymmetries in Population Receptive Field Size and Cortical Magnification Factor in Early Visual Cortex. NeuroImage, 167, 41–52. 10.1016/j.neuroimage.2017.11.021

Slotnick, S. D., Klein, S. A., Carney, T., & Sutter, E. E. (2001). Electrophysiological estimate of human cortical magnification. Clinical Neurophysiology, 112 (7), 1349–1356. 10.1016/S1388-2457(01)00561-2

Talbot, S. A., & Marshall, W. H. (1941). Physiological Studies on Neural Mechanisms of Visual Localization and Discrimination*. American Journal of Ophthalmology, 24 (11), 1255–1264. 10.1016/S0002-9394(41)91363-6

Trimble, C. R. (1968). What Is Signal Averaging. Hewlett-Packard Journal, 19 (8), 2–7.

Van Essen, D. C., Newsome, W. T., & Maunsell, J. H. (1984). The Visual Field Representation in Striate Cortex of the Macaque Monkey: Asymmetries, Anisotropies, and Individual Variability. Vision Research, 24 (5), 429–448. 10.1016/0042-6989(84)90041-5

Varlet, M., Nozaradan, S., Schmidt, R. C., & Keller, P. E. (2023). Neural Tracking of Visual Periodic Motion. European Journal of Neuroscience, n/a(n/a), 1–17. 10.1111/ejn.15934

Wandell, B. A., Dumoulin, S. O., & Brewer, A. A. (2007). Visual Field Maps in Human Cortex. Neuron, 56 (2), 366–383. 10.1016/j.neuron.2007.10.012

